# Increased abundance of Cbl E3 ligases alters PDGFR signaling in recessive dystrophic epidermolysis bullosa

**DOI:** 10.1101/2021.05.10.443425

**Authors:** Esther Martínez-Martínez, Regine Tölle, Julia Donauer, Christine Gretzmeier, Leena Bruckner-Tuderman, Jörn Dengjel

## Abstract

In recessive dystrophic epidermolysis bullosa (RDEB), loss of collagen VII, the main component of anchoring fibrils critical for epidermal-dermal cohesion, affects several intracellular signaling pathways and leads to impaired wound healing and fibrosis. In skin fibroblasts, wound healing is also affected by platelet-derived growth factor receptor (PDGFR) signaling. To study a potential effect of loss of collagen VII on PDGFR signaling we performed unbiased disease phosphoproteomics. Whereas RDEB fibroblasts exhibited an overall weaker response to PDGF, Cbl E3 ubiquitin ligases, negative regulators of growth factor signaling, were stronger phosphorylated. This increase in phosphorylation was linked to higher Cbl protein levels due to increased TGFβ signaling in RDEB. In turn, increased Cbl levels led to increased PDGFR ubiquitination, internalization, and degradation negatively affecting MAPK and AKT downstream signaling pathways. Thus, our results indicate that elevated TGFβ signaling leads to an attenuated response to growth factors, which contribute to impaired dermal wound healing in RDEB.

## Introduction

Epidermolysis bullosa (EB) comprises a group of rare inherited skin disorders characterized by mechanically induced blistering and erosions within the skin and mucosal membranes [1]. Among this group, dystrophic epidermolysis bullosa (DEB) is a monogenetic disorder associated with pathogenic variants in the gene encoding collagen VII (*COL7A1*) [2–4], the main component of anchoring fibrils, which are essential for epidermal-dermal cohesion [5]. The most severe form of the disease is recessive DEB (RDEB). Affected individuals suffer from diminished epidermal-dermal adhesion, skin blistering and subsequent scarring. The continuous damage to the epidermal barrier leads to chronic inflammation and skin infections, which compromise the healing process [6], the result being chronic wounds, progressive scar formation and generalized soft tissue fibrosis. In turn, the inflammatory and fibrotic skin environment favors the development of further disease complications, including squamous cell carcinomas at young ages [7,8]. Despite diverse therapeutic procedures that have been tested, such as partly restoring collagen VII (C7) levels, a cure for the disease does not exist and a high unmet medical need persists [9–12]. Promoting proper healing of skin wounds could be fundamental for improved quality of life and prolonged life expectancy of individuals with RDEB. However, to develop effective regenerative therapies, a deeper understanding of the molecular and cellular mechanisms altered in RDEB is absolutely needed.

During the healing process, one of the first growth factors deposited at the wound site by degranulating platelets is platelet-derived growth factor (PDGF), a major mitogen and stimulator of fibroblasts promoting dermal wound healing [13]. PDGF exists as dimer, composed of A and B chains, forming the homodimers AA and BB and the heterodimer AB [14]. PDGF exerts its action through two specific receptor tyrosine kinases (RTKs), platelet-derived growth factor receptor α and β [15]. Ligand binding results in receptor dimerization and autophosphorylation, which triggers signaling and finally a cellular response. PDGFRα binds all three PDGF isoforms, whereas PDGFRβ preferentially binds PDGF-BB. Although both receptors activate similar signaling pathways and evoke mitogenic signals, PDGFRβ is the more potent in transducing cell motility [14]. For the termination of PDGF signaling, the ligand – receptor complex is internalized via clathrin-coated pits into endosomes, promoted by prior ubiquitination of the receptor molecules by members of the Cbl family of ubiquitin E3 ligases. Most of the receptors are degraded in lysosomes, however, recycling of the receptors can also be triggered under certain circumstances [16–19]. Dysregulation of PDGFR signaling is associated with several malignant processes, such as fibrosis and cancer development in concert with altered TGFβ signaling [20–22] or with the impairment of the wound healing process [23]. Although some features observed as part of the RDEB phenotype might be influenced by PDGFR, little is known about the role of PDGFR signaling in the pathogenesis of RDEB. In this study, we investigate whether the loss of C7 causes the dysregulation of PDGFRβ-mediated signaling in RDEB skin fibroblasts and its potential impact on their wound healing capacity in the dermis.

## Results

### Altered PDGFR-β signaling in RDEB fibroblasts

To study whether loss of C7 could lead to alterations in PDGFRβ signaling, we investigated tyrosine phosphorylation upon stimulation with PDGF-BB in primary normal human fibroblasts (NHF) and RDEB fibroblasts. Three different RDEB fibroblasts and three different NHFs were used for this analysis (see Experimental Procedures for details). After stimulation with PDGF-BB for 0, 5 and 10 min, we observed that RDEB fibroblasts showed in general lower intensities of tyrosine phosphorylation compared to NHF (**Fig. 1A**), indicating an attenuation of PDGFR activity in RDEB.

**Figure 1.**
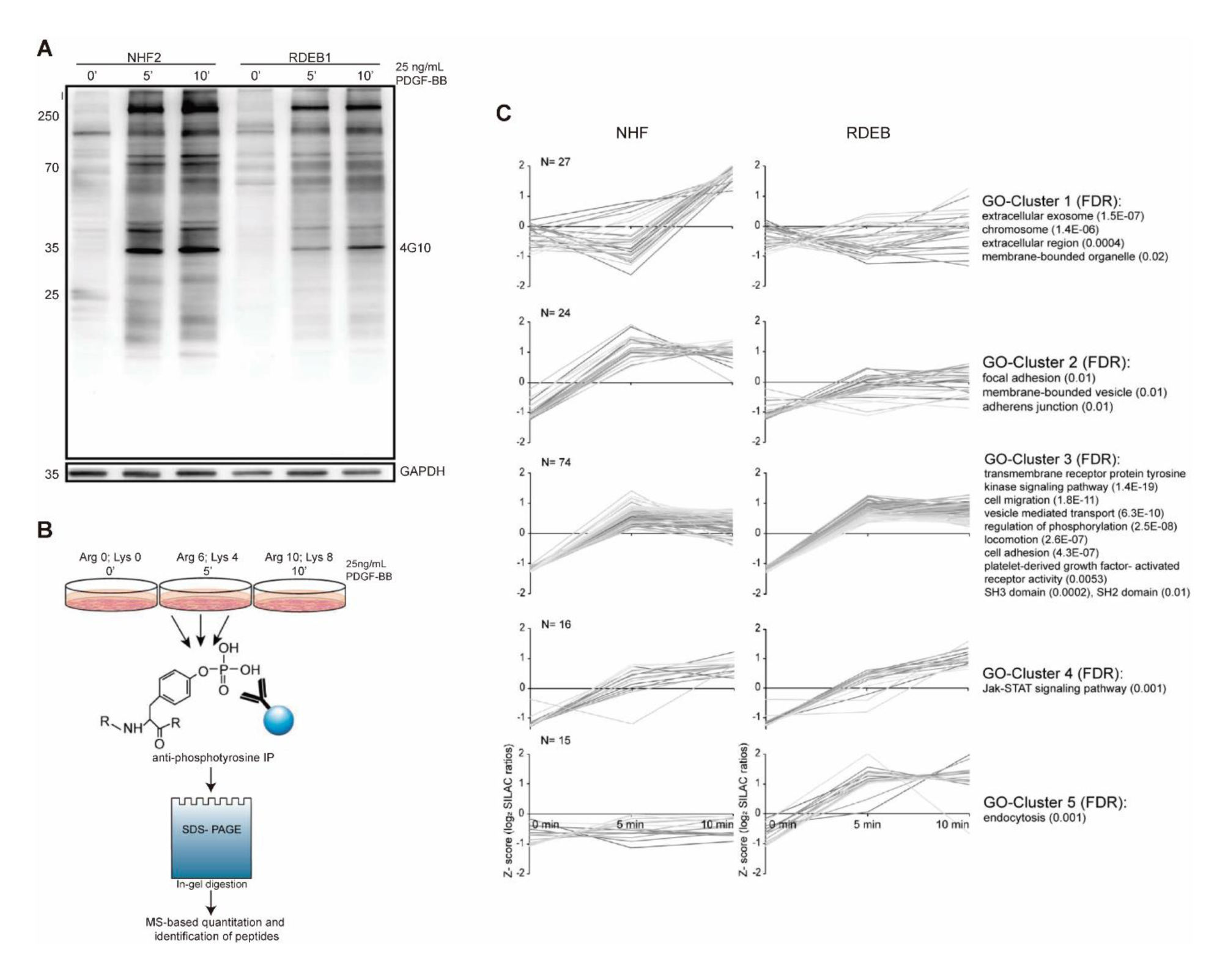
RDEB fibroblasts present differential PDGFR signaling upon stimulation. (**A**) Western blot analysis of phosphotyrosine containing proteins in NHF and RDEB fibroblasts after stimulation with 25 ng/mL PDGF-BB for 5 and 10 min. (**B**) MS-based workflow of the quantitative analysis of PDGFR-based phosphotyrosine signaling. Each cell type was subjected to three different SILAC labels and stimulated with PDGF-BB. Unstimulated cells were used as controls. Three different NHF (NHF2, 4, 8) and RDEB (RDEB1, 4, 5) cells were analyzed. Anti-phosphotyrosine immunoprecipitated samples were in-gel digested and analyzed by LC-MS/MS. (**C**) Cluster analysis of significantly enriched proteins after phosphotyrosine-IP. MS-data of significantly enriched proteins (BH-corrected FDR≤0.05) was *z*-normalized and *k*-means clustered: protein profiles of NHF are shown on the left and RDEB fibroblasts on the right. GO-term enrichment of the proteins in each cluster was computed with a FDR ≤ 0.05 (respective *p*-values are noted in brackets) and a minimum of four proteins per GO-term had to be annotated.

Next, to identify proteins being differentially phosphorylated after PDGF-BB stimulation, we quantified phosphotyrosine-containing proteins by stable isotope labeling by amino acids in cell culture (SILAC)-based mass spectrometry (MS) analyses (**Fig. 1B**) (see Experimental Procedures). Biological replicates with SILAC-label swaps were analyzed to exclude unspecific enriched proteins, contaminants and false-positive hits. In three biological replicates each, we identified 2’324 proteins in total. 156 proteins with a significant enrichment in at least one timepoint were used for the follow-up analysis (Significance A, FDR≤0.05, BH corrected, **Supplementary Table S1**). For the significantly enriched proteins, the average ratios of biological replicates were z-normalized (ungrouped), and subjected to a *k*-means cluster analysis [24]. For NHF and RDEB each cluster contained the same set of proteins, allowing a direct comparison of the cell types (**Fig. 1C**). Clusters 3 and 4 comprised proteins with a similar induction in tyrosine phosphorylation in NHF and RDEB fibroblasts. This cluster was associated with transmembrane receptor protein tyrosine-kinase signaling pathways and platelet-derived growth factor-activated receptor as indicated by GO term analysis (Fisher’s exact test, p<0.05, BH-corrected). SILAC ratios of enriched PDGFRα and PDGFRβ revealed similar inductions in tyrosine phosphorylation in NHF and RDEB fibroblasts, the latter responding slightly weaker (**Supplementary Table S1**). Cluster 1 and 2 contained proteins that were early (Cluster 2) or late (Cluster 1) induced in NHF upon PDGF-BB stimulation, but showed a reduced tyrosine phosphorylation induction in RDEB. These groups contained proteins associated with extracellular exosome, membrane-bounded organelles (Cluster 1) as well as focal adhesion and adherens junction (Cluster 2). Interestingly, cluster 5, which is associated with endocytosis, showed a clear induction of tyrosine phosphorylation after 5 and 10 min stimulation in RDEB fibroblasts, but no induction in NHF.

### Levels of Cbl E3 ligases are increased in RDEB fibroblasts

Cluster 5 contained two interesting proteins that can be linked to altered receptor tyrosine kinase (RTK) activity: c-Cbl and Cbl-b (**Fig. 1C**). They are E3 RING ubiquitin ligases linked to ubiquitination and degradation of several RTKs upon growth factor stimulation, PDGFRβ being one of them [18]. In agreement with the cluster analysis, c-Cbl and Cbl-b were 14 and 2 times higher enriched, respectively, in RDEB fibroblasts than in NHF after 5 min of PDGF-BB stimulation (**Fig. 2A**). Thus, phosphorylated levels of these proteins were higher in RDEB fibroblasts upon growth factor stimulation, which points to an elevated activity of them [17,18]. Also by western blot analysis, c-Cbl and Cbl-b were more enriched in phosphotyrosine immunoprecipitations in RDEB fibroblasts stimulated by PDGF-BB (**Fig. 2B**), confirming that RDEB fibroblasts present increased phosphorylated levels of c-Cbl and Cbl-b after PDGFR stimulation.

**Figure 2.**
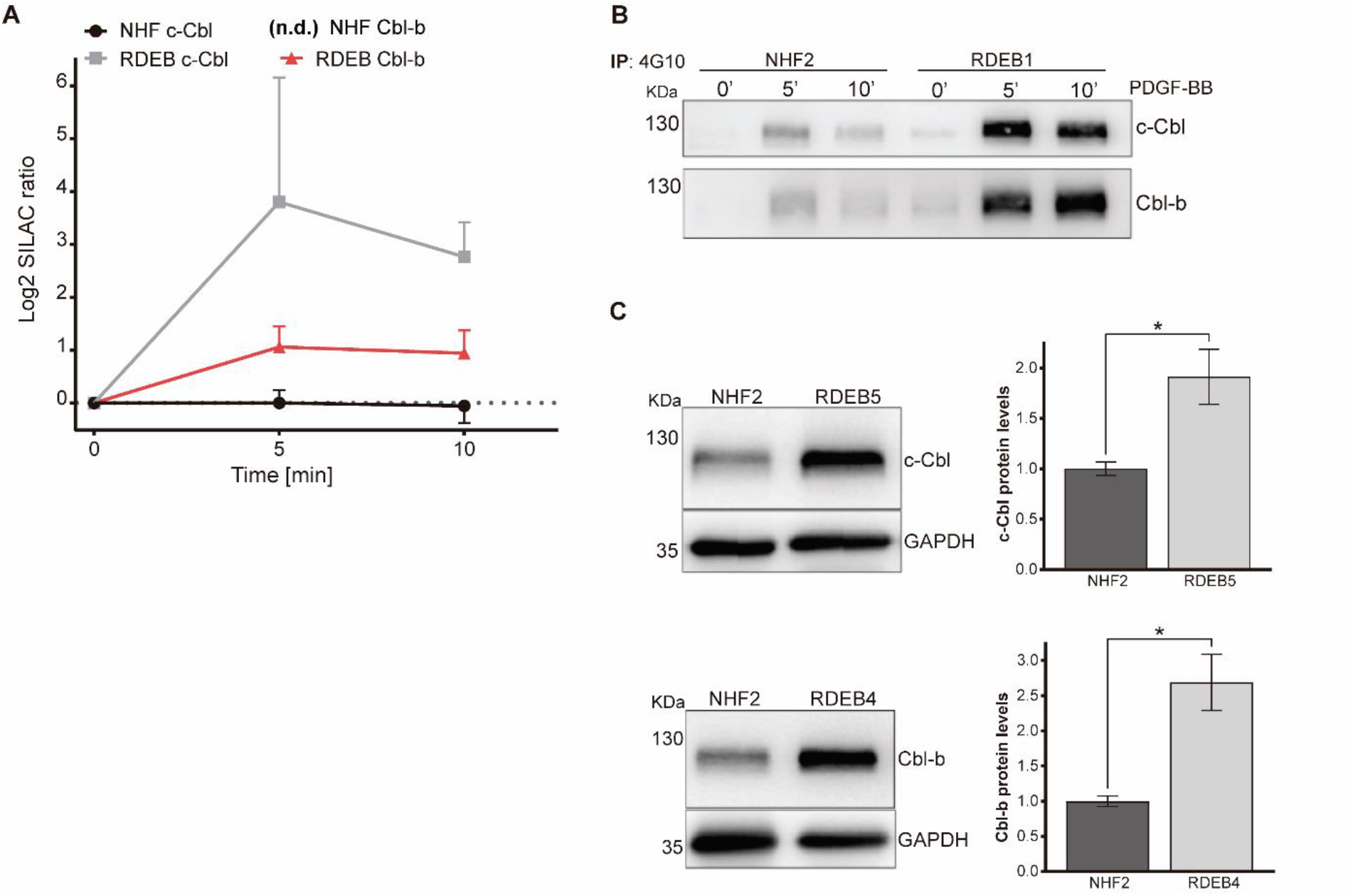
Expression of Cbl E3 ubiquitin ligases is increased in RDEB fibroblasts. (**A**) Log_2_SILAC ratios of c-Cbl and Cbl-b, proteins identified in cluster 5, extracted from the cluster analysis obtained from LC-MS/MS experiments represented in Fig. 1C. n.d. not detected. (**B**) Representative western blot of Cbl-b and c-Cbl after anti-phosphotyrosine IP of NHF and RDEB fibroblasts stimulated with 25 ng/mL PDGF-BB for the indicated timepoints. (**C**) Western blot and quantification of Cbl-b and c-Cbl levels in whole cell lysates of NHF and RDEB fibroblasts. Data are represented as mean ± s.e. (n=3), * *p* ≤ 0.05 using Student’s t-test for the comparison between NHF and RDEB fibroblasts.

To test whether the increase of c-Cbl and Cbl-b observed by LC-MS/MS and western blot is due to an increase in their phosphorylation or their abundance, we analyzed whole cell lysates of RDEB fibroblasts and NHF. Interestingly, RDEB fibroblasts presented higher protein levels of Cbl-b and c-Cbl (**Fig. 2C**), indicating that the higher phosphorylation levels are due to a dysregulation of Cbl protein abundance and not PDGFR activity. This agrees to the observation that PDGFR itself was not substantially dysregulated in RDEB fibroblasts compared to NHF (**Supplementary Table S1**). Altogether, these results indicate that RDEB fibroblasts present elevated levels of c-Cbl and Cbl-b that, upon PDGF-BB stimulation, become activated through phosphorylation, which might affect PDGFRβ activity in RDEB fibroblasts.

### Rapid internalization and degradation of PDGFRβ upon stimulation in RDEB fibroblasts

Cbl E3 ubiquitin ligases are implicated in negative regulation of RTKs by inducing their ubiquitination and subsequent degradation [18]. To study if elevated levels of c-Cbl and Cbl-b in RDEB fibroblasts affect the half-life of PDGFRβ, fibroblasts were stimulated with PDGF-BB for different timepoints and PDGFRβ levels were analyzed. We found that 5 min treatment with PDGF-BB led to practically complete degradation of the receptor in RDEB fibroblasts, while in NHF we only observed approximately 50% degradation (**Fig. 3A**). Since PDGFRβ ubiquitination by the aforementioned E3 ligases is the signal for degradation, we investigated ubiquitination levels of the receptor upon growth factor stimulation. NHF and RDEB fibroblasts were subjected to a 3 min treatment with 15 ng/mL PDGF-BB and PDGFRβ was immunoprecipitated for analyzing ubiquitination levels. Indeed, RDEB fibroblasts presented up to 4-times higher ubiquitination levels of PDGFRβ than NHF (**Fig. 3B**). Moreover, we again observed a rapid degradation of PDGFRβ in RDEB fibroblasts, since PDGFRβ levels were reduced upon 3 min stimulation, while in NHFs there was no reduction (**Fig. 3B**, left bottom panel).

**Figure 3.**
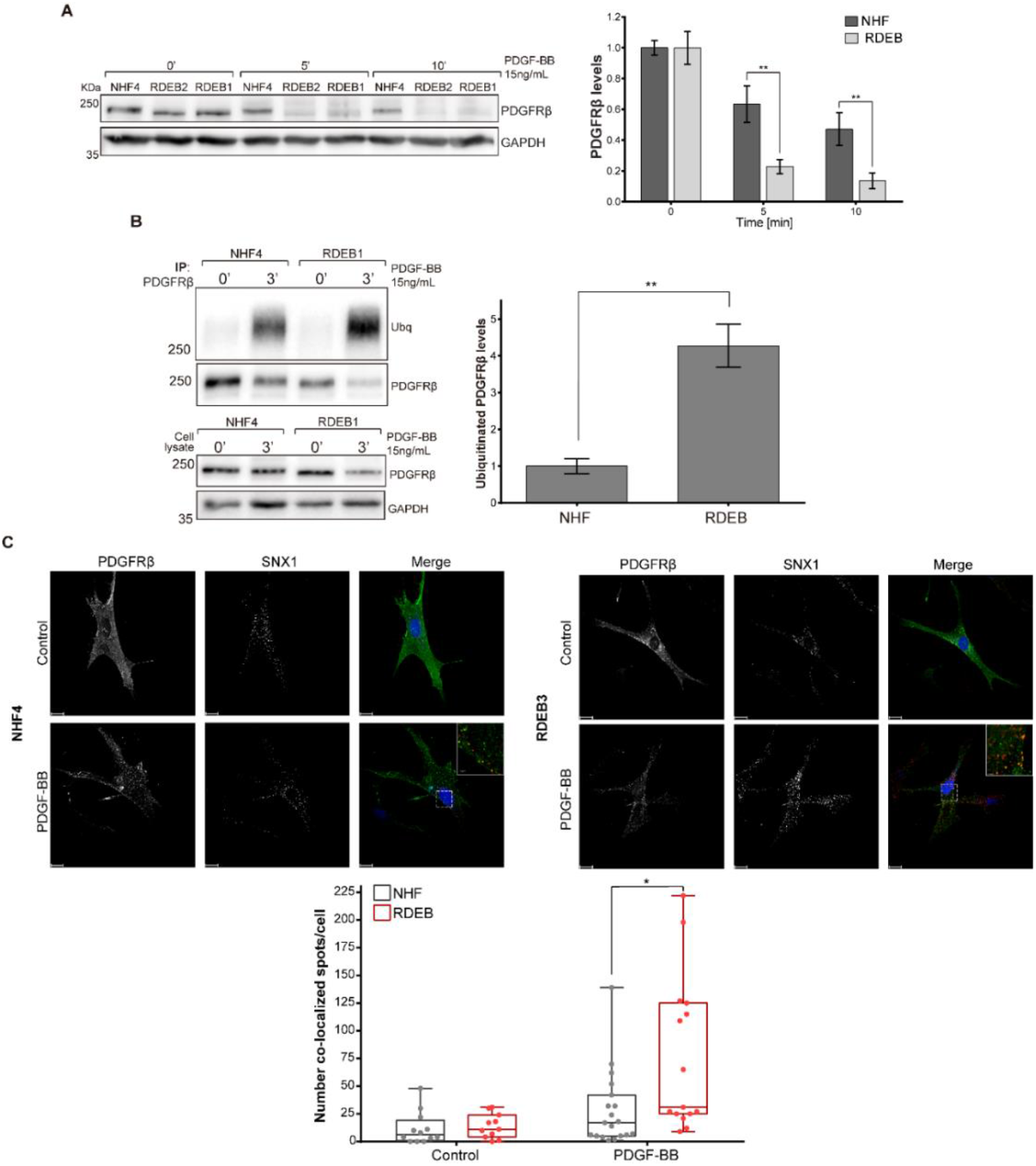
Increased PDGFRβ degradation upon growth factor stimulation in RDEB fibroblasts. (**A**) Representative western blot of PDGFRβ and Cbl-b levels (left panel) and quantification of PDGFRβ (right panel) in NHF and RDEB fibroblasts in control conditions and after stimulation with 15 ng/mL PDGF-BB for the indicated timepoints. Data was normalized to GAPDH levels. Comparisons were performed between NHF and RDEB cells at the respective timepoints. Data are represented as mean ± s.e (n=3), including two different NHF (NHF4, 9) and three RDEB (RDEB1-3) types in each experiment. ** *p* ≤ 0.01 using Student’s t-test computed with a FDR ≤ 0.05 for multiple testing. (**B**) Representative western blot (left panel) and quantification (right panel) of PDGFRβ ubiquitination after 3 min of stimulation with 15 ng/mL PDGF-BB. PDGFRβ was immunoprecipitated prior ubiquitin detection. Corresponding whole cell lysates were analyzed. Ubiquitinated levels were normalized to the corresponding levels of immunoprecipitated PDGFRβ. Data are represented as mean ± s.e (n=3), including two different NHF and three RDEB types in each experiment. ** *p* ≤ 0.01 using Student’s t-test for the comparison between NHF and RDEB fibroblasts. (**C**) Representative image (upper panel) and quantification (bottom panel) of immunofluorescence analysis of PDGFRβ/SNX1 co-localization. NHF and RDEB cells were stimulated 5 min with 15 ng/mL PDGF-BB. Cells were co-immunostained with specific anti-PDGFRβ (green) and anti-SNX1 (red) antibodies and analysed by confocal microscopy. Nuclei were stained with Hoechst 33258 (blue). Scale bars, 15 μm. Insets = higher magnification of the relative areas within the white-dotted boxes. Scale bars, 2 µm. Quantification was performed by object-based co-localization extension with Imaris software. Data represented in box plot showing the number of co-localized PDGFRβ/SNX1 spots per cell in each condition. Between 15 and 19 cells were counted per condition, each of them including two NHF (NHF4, 9) and two RDEB (RDEB 2-3) cell types. * *p* ≤ 0.05 using Student’s t-test computed with a FDR ≤ 0.05 for multiple testing.

To study PDGFRβ internalization upon PDGF-BB stimulation, we also performed immunofluorescence microscopy and analyzed its co-localization with sortin nexin-1 (SNX1), an endosomal protein described to play a role in sorting RTKs to lysosomes for their degradation upon growth factor stimulation [25,26]. NHF and RDEB fibroblasts were treated for 5 min with 15 ng/mL PDGF-BB and co-immunostained for PDGFRβ and SNX1. Upon stimulation, the receptor was internalized in both cell types changing to a more punctuate, cytoplasmic pattern (**Fig. 3C**, PDGFRβ panels). Likewise, SNX1 puncta presence increased in the cytoplasm of PDGF-treated cells, being more intense in RDEB fibroblasts (**Fig. 3C**, SNX1 panels). Co-localization between SNX1 and PDGFRβ was observed in both cell types, indicating that receptor molecules were directed to lysosomes for degradation, but to a significantly greater extent in RDEB (**Fig. 3C**, Merge panel and bottom panel), indicating that the degradation process is faster in those cells. Collectively, these results indicate that, upon PDGF-BB stimulation, PDGFRβ is more rapidly ubiquitinated, internalized and degraded in RDEB fibroblasts likely due to the presence of higher levels of Cbl E3 ligases.

### RDEB fibroblasts present an attenuated response to PDGF-BB stimulation

To study whether faster PDGFRβ degradation in RDEB fibroblasts influences its downstream signaling, we investigated the effect in two central pathways regulated by PDGFRβ: the mitogen-activated protein kinase/extracellular signal-regulated kinases (MAPK/ERK) pathway [27,28] and the phosphatidylinositol 3-kinase (PI3K)/AKT pathway [29,30]. NHF and RDEB fibroblasts were subjected to up to 8 h of PDGF-BB stimulation and phosphorylated ERK1/2 and AKT levels were analyzed. We found that in both cell types PDGFRβ activation led to the phosphorylation of ERK1/2 and AKT, but in NHF the signal was more stable and lasted longer than in RDEB (**Fig. 4A**).

**Figure 4.**
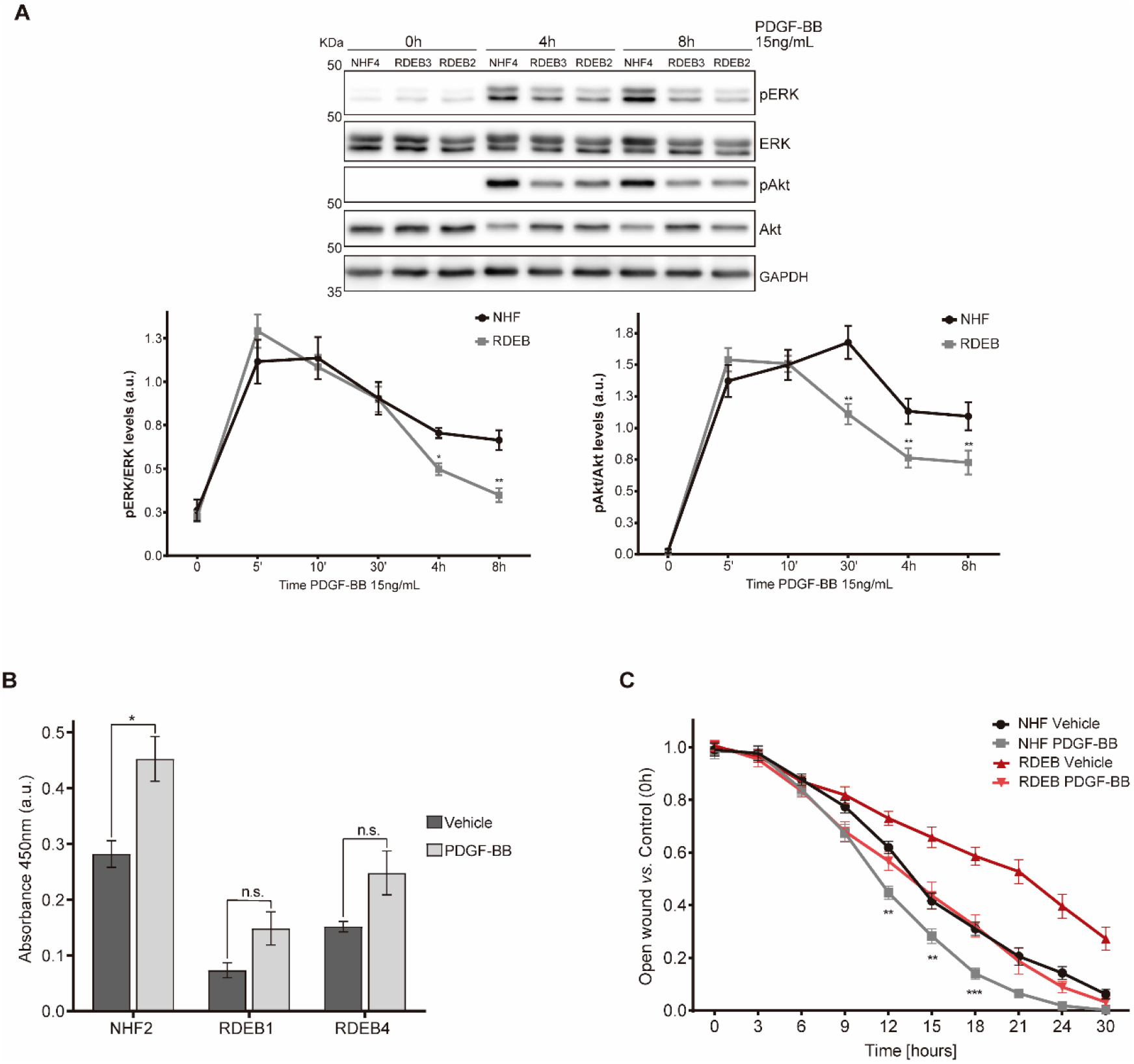
Attenuated PDGF-induced MAPK signaling in RDEB fibroblasts compared to NHF. (**A**) Representative western blot (upper panel) and quantification (bottom panel) of phosphorylated ERK1/2 (pERK) and Akt (pAkt) levels after stimulation with 15 ng/mL PDGF-BB at the indicated timepoints. Data were normalized to GAPDH and total ERK1/2 (pERK/ERK) or Akt (pAkt/Akt). Data are represented as mean ± s.e (n=3) each experiment including two different NHF (NHF4, 9) and three RDEB (RDEB1-3) cell types. * *p* ≤ 0.05, ** *p* ≤ 0.01 using Student’s t-test computed with a FDR ≤ 0.05 for multiple testing. (**B**) BrdU cell proliferation assay was performed in NHF and RDEB fibroblasts treated with vehicle or 25 ng/mL PDGF-BB for 24 h. The assay was done under low serum conditions (0.25% FBS). Data are represented as mean ± s.e. (n=3), * *p* ≤ 0.05, n.s. = non-significant using a two-way ANOVA with a Holm-Sidak test post hoc analysis. Note: values of NHF and RDEB cells were significantly different in all conditions, *p* ≤ 0.05. (**C**) NHF and RDEB fibroblasts were subjected to scratch assays in low-serum conditions (0.25% FBS) in presence of 1 µg/mL of mitomycin C. Cells remained untreated (vehicle) or treated with 5 ng/mL PDGF-BB. The motility of cells was monitored by time - lapse video microscopy (Incucyte® Live Cell analysis system, Essen Biosciences) and wound closure quantification was performed with ImageJ (NIH) software. NHF and RDEB data was normalized to their respective control (0 h). Data are represented as mean ± s.e. (n=3) each experiment including two different NHF (NHF4, 9) and RDEB (RDEB1, 2) cells. Statistical comparisons were made between NHF and RDEB stimulated with PDGF-BB at the respective timepoints. ** *p* ≤ 0.01 and \*\*\*\**p* ≤ 0.0001 using two-way ANOVA with a Tukey test post hoc analysis.

It is known that MAPK and PI3K/AKT signaling regulate different cellular processes, such as proliferation or cell motility, which are important in the wound healing processes [27,30]. Given our previous results and the described impairment of the wound healing process in RDEB [31], we investigated cell proliferation upon PDGF-BB stimulation. RDEB fibroblasts showed significantly diminished proliferation compared to NHF, both in basal conditions and after growth factor stimulation (**Fig. 4B**). Importantly, whereas NHF showed a significant increase in cell proliferation after PDGF-BB stimulation, RDEB fibroblasts responded weaker, in a non-significant manner. Next, we performed a scratch assay for studying cell motility. After inflicting the scratch, NHF and RDEB fibroblasts were treated with vehicle or PDGF-BB and cell migration was followed for 30 h. In order to block cell proliferation, fibroblasts were treated with mitomycin C in all assayed conditions. We found that cell motility was also affected in RDEB fibroblasts both in basal conditions and after PDGF-BB stimulation. Although the migration rate increased in response to PDGF-BB, RDEB fibroblasts never reached the migration levels of NHFs (**Fig. 4C**). Taken together, these results indicate that RDEB fibroblasts are capable of responding to PDGF-BB stimulation; however, their response is impaired compared to NHFs, with different kinetics of ERK and AKT activation and lower proliferati ve and migratory capacities.

### Altered TGFβ signaling in RDEB fibroblasts induces elevated Cbl levels

In RDEB, increased TGFβ receptor signaling is considered as one of the main pathways implicated in impairment of the wound healing process and in maintaining a characteristic, inflammatory, fibrotic microenvironment [32–34]. As TGFβ stimulation was shown amongst others to lead to an increase in c-Cbl expression in mouse fibroblasts [35], we tested if increased TGFβ levels might be the cause for elevated c-Cbl and Cbl-b protein levels in RDEB fibroblasts. We verified that Cbl-b levels were increased in a time-dependent manner upon TGFβ stimulation in NHF (**Fig. 5A**), suggesting that elevated levels of c-Cbl and Cbl-b in RDEB fibroblasts could be caused by the upregulation of TGFβ signaling. Phosphorylation of Smad3, a well-known target of the TGFβ pathway, was used as a readout for an effective activation of the receptor (**Fig. 5A**). Next, TGFβ signaling was blocked in RDEB fibroblasts with LY210976, a TGF-β receptor type I/II (TβRI/II) dual inhibitor. Indeed, Cbl-b levels were downregulated after 24 h of LY210976 treatment, reaching similar levels as observed in NHF (**Fig. 5B**). Again, phosphorylated Smad3 was used as readout of effective blockade of the receptor. Collectively, these results indicate that elevated levels of c-Cbl and Cbl-b in RDEB fibroblasts are caused by the upregulation of TGFβ signaling, and that blocking this signal leads to a decrease in the levels of the E3 ubiquitin ligases (**Fig. 6**).

**Figure 5.**
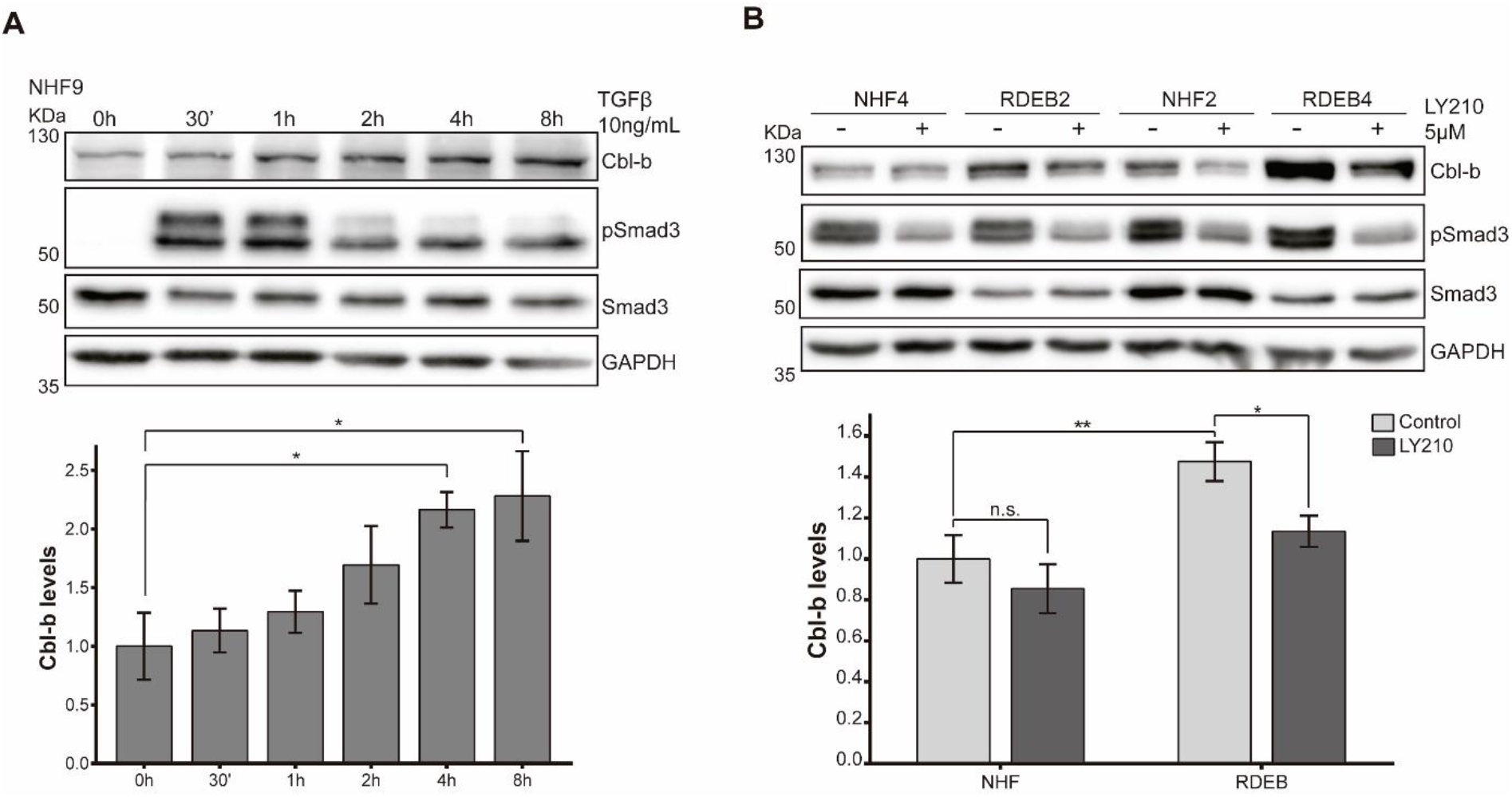
TGFβ regulates the differential expression of Cbl E3 ubiquitin ligases in RDEB fibroblasts. (**A**) Representative western blot of Cbl-b and phosphorylated Smad3 (pSmad3) levels and quantification of Cbl-b levels in NHF subjected to 10 ng/mL TGFβ treatment at different timepoints. Data was normalized to GAPDH. Cbl-b levels were compared with the control group (0 h). Data are represented as mean ± s.e (n=3) each experiment including two different NHF (NHF4, 9). Statistical comparison was made with respect the control (0 h). * *p* ≤ 0.05 using one-way ANOVA with a Dunnet test post hoc analysis. (**B**) Representative western blot and quantification of Cbl-b levels upon TGFβ signaling blockade with LY210976 (LY210) 5 µM for 24 h. Data was normalized to GAPDH levels and are represented as mean ± s.e (n=3) including three different NHF (NHF2, 4, 9) and three RDEB (RDEB2, 3, 4) cell types. * *p* ≤ 0.05 and \*\**p* ≤ 0.01 using Student’s t-test. n.s. = non-significant.

**Figure 6.**
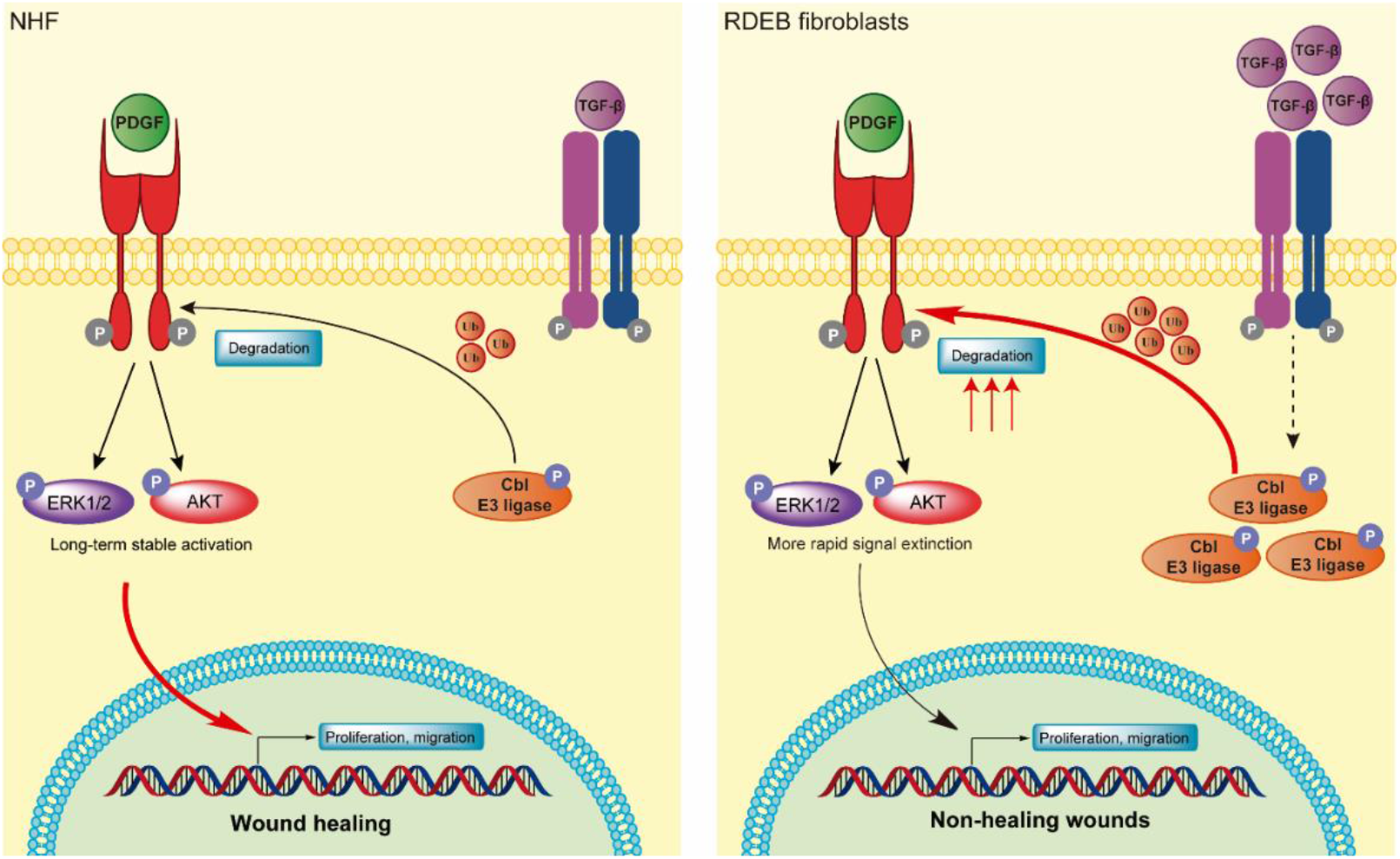
Proposed mechanisms. In RDEB, increased TGFβ signaling compared to control conditions induces an increase in the levels of E3 ubiquitin ligases c-Cbl and Cbl-b. This leads to a more rapid ubiquitination, internalization and degradation PDGFRβ upon PDGF-BB stimulation, which in turn provokes a more rapid extinction of the downstream signaling and, subsequently, limited cellular responses, such as proliferation and cell motility, key processes for proper wound healing.

## Discussion

PDGFRβ signaling is crucial for fibroblast homeostasis and regulates diverse processes such as wound healing and fibrosis [20,36–38], both of which are dysregulated in RDEB [6,7,31]. However, little is known about PDGFR signaling in RDEB and its potential role in wound healing and fibrosis upon loss of C7. Here, we present data that elevated TGFβ signaling due to the loss of C7 affects PDGFRβ signaling and how this is translated in the impairment of processes related to wound healing, i.e. cell proliferation and migration. Employing an unbiased phosphoproteomics approach, we observed overall attenuated PDGFRβ signaling upon PDGF-BB treatment in RDEB fibroblasts. In total 51 proteins responded in NHF but not in RDEB fibroblasts to PDGF-BB stimulation. Respective proteins were associated with GO terms related to the extracellular region and molecular mechanisms associated with the cell membrane, such as ECM-receptor interaction, focal adhesions and membrane-bound vesicles, in accordance with the fact that RDEB fibroblasts present alterations in the ECM, which provokes changes in cell-ECM interactions [34]. Of note, we found two members of the Cbl family of RING finger-containing E3 ubiquitin ligase, c-Cbl and Cbl-b, to not follow this trend and to be increasingly phosphorylated in RDEB fibroblasts.

Cbl E3 ligases are important negative regulators of RTKs. They are activated by tyrosine phosphorylation after growth factor stimulation and induce ubiquitination, internalization and degradation of RTKs [17,18]. Our data show that RDEB fibroblasts not only present higher levels of phosphorylated c-Cbl and Cbl-b upon PDGF-BB stimulation compared to NHF, but that they have increased overall levels of these proteins. Therefore, one might predict that attenuated PDGFRβ signaling in RDEB fibroblasts could be caused by enhanced levels of Cbl E3 ligases, which provoke a prompt downregulation of the receptor upon PDGF-BB stimulation, leading to reduced signal transduction. Our results showed that, upon PDGF-BB stimulation, PDGFRβ is more rapidly ubiquitinated, internalized and degraded in RDEB compared to normal fibroblasts. Analysis of ERK1/2 and AKT phosphorylation, two of the main pathways regulated by PDGFRβ related to mitogenic signaling and cell motility [27,29], revealed that in RDEB fibroblasts both pathways are only transiently activated compared to NHF upon PDGF-BB stimulation. This result verified that the more rapid degradation of PDGFRβ in RDEB after its activation leads to reduced signal transduction.

To explore whether the differences in ERK and AKT signaling led to differences in cell responses, we analyzed cell proliferation and motility in PDGF-BB stimulated fibroblasts. Both processes are tightly related with wound healing. As expected, RDEB fibroblasts presented less proliferative and migratory capacities in basal conditions when compared to NHF. Nonetheless, RDEB fibroblasts were capable of responding to PDGF-BB, since we observed an increase in proliferation and migration rates after stimulation. However, their proliferative and migratory capacities were always lower than the ones of NHF. These results reinforce the hypothesis that the response triggered in RDEB fibroblasts after PDGFRβ activation is abnormal.

Since it was reported that increased TGFβ activity induced the expression of c-Cbl in mice [35], and TGFβ signaling is upregulated in RDEB, being one of the most important modulators of this disease [31,34,39], we investigated whether TGFβ upregulation might induce the increase in Cbl protein levels. Our data support this hypothesis, since activation of TGFβR in NHF led to an increase of Cbl-b levels in a time-dependent manner. Moreover, blocking TGFβ signaling with a pharmacological inhibitor in RDEB fibroblasts led to a decrease in Cbl-b levels, strengthening the aforementioned hypothesis. The mechanism by which TGFβ induces Cbl expression remains to be determined and is an active topic of our current investigations.

Taken together, our data show that increased TGFβ signaling occurring in RDEB fibroblasts induces an increase in the levels of E3 ubiquitin ligases c-Cbl and Cbl-b. This leads to attenuated PDGFRβ signaling due to a more rapid internalization and degradation upon PDGF-BB stimulation, which in turn provokes a more rapid extinction of downstream signaling and, subsequently, limited cellular responses, such as proliferation and cell motility, key processes for proper wound healing (**Fig. 6**). Loss of C7 induces numerous alterations in ECM and cell homeostasis apart from the explored ones, known and unknown, that may act as disease modifiers and might contribute to aberrant ERK signaling. Thus, further work is necessary to fully understand the role of PDGFRβ and other disease modifiers in the dynamic processes of healing and fibrosis in RDEB. This study provides new understanding of the molecular and cellular pathogenesis of RDEB and consequently new insights into potential therapy options. In the clinic, PDGF-BB has been topically applied to enhance the healing of chronic wounds and loss of myofibroblast features in a MAPK dependent manner [40,41]. However, in the light of our findings, this treatment might not be efficient in RDEB due to chronically increased activation of TGFβ signaling.

## Experimental Procedures

### Primary skin cells culture

Normal human primary fibroblasts (NHF) were isolated from the foreskin of circumcised 2 -, 4-, 8- and 9-year old healthy boys. Primary dermal fibroblasts from RDEB patients were obtained from skin specimens of 6 different donors who were less than one year of age [34,42]. RDEB fibroblasts used in this study were selected based on the absence of C7 as detected by Western blotting. The study was approved by the Ethics Committee of Freiburg University (# 45/03 - 110631) and conducted according to the Declaration of Helsinki.

Fibroblasts were cultured in DMEM with 4,5 g/L glucose, L-glutamine and sodium pyruvate (PAN-Biotech, Aidenbach, Germany) supplemented with 10% fetal bovine serum (FBS) (ThermoFisher Scientific, Waltham, MA) and 1% Penicillin/Streptomycin (PAN-Biotech, Aidenbach, Germany) and maintained at 37°C in a 5% CO2-humidified atmosphere. Cells were harvested with trypsin/EDTA solution (0.05%/0.02% weight/volume) (PAN-Biotech, Aidenbach, Germany). For quantitative analysis of proteins by mass-spectrometry, fibroblasts were SILAC-labeled. Cells were cultured in high-glucose SILAC-DMEM (Thermo Fisher Scientific) supplemented with 10% dialyzed fetal bovine serum (ThermoFisher Scientific, Waltham, MA), 1% Glutamax supplement (ThermoFisher Scientific), 1% Penicillin/ Streptomycin (PAN-Biotech) and 42 mg/L of L-arginine and 73 mg/L of L-lysine (Sigma). To reduce the arginine-to-proline conversion, 82 mg/L of L-proline was added to the media. Per experimental setup, the media was either supplemented with L-lysine-^2^H_4_ and L-arginine-^13^C_6_-^14^N_4_ (named as Lys4, Arg6) (Sigma-Aldrich, St. Louis, MO), L-lysine-^13^C_6_-^15^N_2_ and L-arginine-^13^C_6_-^15^N_4_ (named as Lys8, Arg10) (Sigma-Aldrich) or L-lysine and L-arginine (named as Lys0, Arg0) (Sigma-Aldrich). Fibroblasts were maintained in SILAC media for two weeks prior the performance of the experiments to ensure full amino acid labeling.

### Reagents

Recombinant human platelet-derived growth factor-BB (PDGF-BB) and recombinant human transforming Growth Factor-β1 (TGFβ1; HEK293 derived) were purchased from PeproTech (Hamburg, Germany) and they were dissolved in sterile ddH2O. TGF-β receptor inhibitor LY2109761 was purchased from MedChem Express (Monmouth Junction, NJ, USA) and it was resuspended in sterile dimethylsulfoxid (DMSO; Sigma-Aldrich). Primary antibodies anti - PDGFRβ (D-6; sc-374573), anti-ubiquitin (A-5; sc-166553), GAPDH (FL-335; sc-25778) and β-actin (C4; sc-47778) were purchased from Santa Cruz Biotechnologies (Heidelberg, Germany). Anti-phospho-Smad3 (S423 + S425) (EP823Y; ab52903) and anti-Smad3 (ab28379) were purchased from Abcam (Cambridge, UK). Antibodies anti-phospho-ERK 1/2 (T202/Y204) (#9101), anti-ERK 1/2 (#9102), anti-phospho-Akt (Ser473) (#9271), anti-Akt (#9272), PDGFRβ (28E1; #3169), anti-Cbl-b (D3C12; # 9498), anti-c-Cbl (D4E10; # 8447) were purchased from Cell Signaling Technology (St Louis, MO, USA). Anti-phosphotyrosine antibody conjugated to agarose beads (clone 4G10; 16-101) and anti-phosphotyrosine antibody (Clone 4G10; 05-321) were obtained from Merck Millipore (Darmstadt, Germany). Anti-SNX1 (Clone 51; 611482) antibody was purchased from BD Transduction Laboratories (BD Biosciences, Breda, The Netherlands). Horseradish peroxidase (HRP)-conjugate secondary antibodies anti-rabbit (111-035-045), anti-mouse (115-035-062) and anti-mouse light chain specific (115-035-174) were purchased from Jackson ImmunoResearch Laboratories (Newmarket, UK). Alexa Fluor 488 anti-rabbit and Alexa Fluor 594 anti-mouse were purchased from Invitrogen Life Technologies.

### Phosphotyrosine enrichment by immunoprecipitation

Fibroblasts were serum-starved overnight prior to stimulation with 25 ng/mL of PDGF-BB for 0, 5, or 10 min. Cells were lysed in Triton buffer (50 mM Tris-HCl pH 7.4, 1% Triton-X 100, 137 mM NaCl, 1% Glycerine; Sigma-Aldrich) supplemented with phosphatase and protease inhibitors (cOmplete™ Protease Inhibitor Cocktail and PhosSTOP™; Roche, Rotkreuz, Switzerland). Cells were lysed for 2 h on ice. The protein concentration was determined by pierce BCA protein assay kit (ThermoFisher Scientific) and lysates were mixed in a 1:1:1 protein ratio. For the immunoprecipitation of phosphotyrosine residues, lysates were incubated with 75 μg of anti-Phosphotyrosine antibody conjugated to agarose beads (clone 4G10) for 4 h at 4°C. After immunoprecipitation, the beads were washed with lysis buffer and proteins were eluted in 2x Laemmli loading buffer containing 1 mM dithiothreitol (DTT; Amresco, Solon, OH) for 30 min. at 75°C.

### MS-based analysis of phosphotyrosine signaling events

After immunoprecipitation of phosphotyrosine residues as described above, the reduced samples were alkylated with 5.5 mM iodoacetamide (IAA; Sigma-Aldrich). Proteins were separated by 4-12% Bis-Tris gradient gels (NuPAGE, Invitrogen, ThermoFisher Scientific) and protein lanes were divided into 10 fractions. Proteins from the gel pieces were digested with trypsin HPLC-grade (Promega AG, Dübendorf, Switzerland) overnight and the resulting peptide solutions were desalted on C18-based STAGE tips as previously described [43,44]. Samples were analyzed by an LTQ Orbitrap XL mass spectrometer (ThermoFisher Scientific) coupled to an Agilent 1200 nano flow-HPLC, as well as by a QExactive mass spectrometer (Thermo Fisher Scientific), coupled to an Easy nanoLC (ThermoFisher Scientific). HPLC column tips (fused silica, 75 μm inner diameter, 20 cm final length) were in-house packed with Reprosil-Pur 120 C18-AQ, 1.9 μm (Dr. Maisch). A gradient of buffer A (0.5% acetic acid; Sigma-Aldrich) and B (0.5% acetic acid in 80% acetonitrile; LGC Standards-Promochem, Wesel, Germany) with increasing organic proportion was used for peptide separation. For the LTQ Orbitrap samples were separated within 145 min (linear gradient of buffer B from 10% to 80% within 110 min). On the Easy nanoLC, samples were separated for 50 min (5-25% buffer B in 37 min, increase to 50% at 45 min, and complete elution at 80% buffer B). The flow rates were set to 250 nl.min^-1^. The spray voltage was 2.3 kV on the LTQ Orbitrap XL, 3.9 kV for the QExactive. The capillary temperature was set to 200°C for the LTQ Orbitrap XL and 320°C for the QExactive. No sheath or auxiliary gas flow was used. Data acquisition was done data - dependent: the MS switched automatically between MS (max. 1×106 ions) and a maximum of five MS/MS scans (10 MS/MS scans on the QExactive). The collision energy in the linear ion trap was set to 35% (25% for the QExactive) and a target value of 5’000 parent ions. The scan range for the LTQ Orbitrap XL was set to 350-2000 m/z with a resolution of 60’000. For the QExactive the scan range was set to 200-2000 m/z with a resolution of 70’000. Parent ions with an unassigned charge or a charge state of z= 1 were excluded from fragmentation.

### Protein identification and relative quantification

Peak detection and relative quantification of SILAC-pairs was conducted by MaxQuant version 1.4.1.2 [45]. Peaks were searched against a full-length human Uniprot database from December 2017. Enzyme specificity was assigned to trypsin/P and quantitation was based on three SILAC labels. Carbamidomethylation was set as a fixed modification; N-terminal acetylation and oxidation of methionine, as well as phosphorylations of serine/threonine and tyrosine were set as variable modifications. The MS/MS tolerance was set to 0.5 Da. A maximum of two missed cleavages was allowed. Proteins were identified with a minimum of one unique peptide. For quantification, a ratio count of two was needed. Identification and quantification of peptides and proteins was based on a reverse-database with a false discovery rate (FDR) of 0.01. Peptide length had to be at least 7 amino acids. Identified peptides were re-quantified. For enhanced identification, runs were matched with a match time window of 1 min. For the data analysis, normalized protein ratios were evaluated using Perseus 1.5.5.3 [46]. Contaminants, reverse hits and proteins only identified by side were excluded from further analysis. To eliminate unspecific enriched proteins, Significance A with a Benjamini-Hochberg corrected FDR of 0.05 was calculated for normalized, log2-transformed ratios. Only proteins with a significant enrichment in at least one experiment were used for subsequent analysis. The ratios were z - normalized and k-means clustered in MeV (Multi Experiment viewer) [24]. Enriched GO-terms (FDR = 0.05) in 5 different clusters were identified in STRING version 10.5 [47].

### Western blot analysis

Normal and RDEB-derived fibroblasts were cultured in 6-cm plates or 6-well plates until they reached 80% confluence. Cells were serum-starved overnight prior to stimulation with PDGF - BB or TGFβ. Cells were lysed with modified RIPA buffer (50 mM Tris-HCl pH 7.5, 150 mM NaCl, 1mM EDTA, 1% NP-40, 0.25% deoxycholic acid; Sigma-Aldrich) supplemented with proteases and phosphatases inhibitors. Lysates were centrifuged at 12000 rpm for 15 min at 4°C. Protein concentration was determined by pierce BCA protein assay kit and, when possible, samples containing 30 µg protein were prepared in Laemmli loading buffer containing 1mM DTT. Samples were boiled 10 min at 75°C. Proteins were separated in SDS-PAGE gels and transferred to 0.2 μm PVDF membranes (Amersham™ Hybond™ Protran). The membranes were blocked during 1h in 5% BSA or 5% non-fat milk prepared in TBS with 0.1% Tween-20 (TBST) and they were incubated overnight at 4°C with the following primary antibodies and dilutions: anti-PDGFRβ (D-6) (1:500), anti-Cbl-b (1:1000), anti-c-Cbl (1:1000), anti-phospho-ERK 1/2 (1:2000), anti-ERK 1/2 (1:1000), anti-phospho-Smad3 (1:2000) and anti-GAPDH (1:1000) and anti-β-Actin (1:3000) as loading control. The next day, blots were incubated 1 h at room temperature with goat anti-rabbit (1:10000) or anti-mouse (1:5000) HRP-conjugated secondary antibodies (Jackson ImmunoResearch Europe Ltd, Newmarket, UK) and signal was detected with WesternBright ECL Spray (Advansta, Menlo Park, CA, USA) or SuperSignal West Femto Maximum Sensitivity Substrate (ThermoFisher Scientific) in Odyssey^®^ Fc Imaging System (LI-COR Biosciences, Lincoln, NE). Densitometric analysis for quantification were done with Image-J (NIH) software.

### PDGFRβ ubiquitination levels

Normal and RDEB-derived fibroblasts were cultured in 10-cm plates until they reached 90% confluence. Cells were serum-starved overnight prior to 3 min stimulation with 15 ng/mL PDGF-BB. Cells were lysed for 30 min at 4°C on the wheel with RIPA buffer (50 mM Tris-HCl pH 8.0, 150 mM NaCl, 1% NP-40, 0.5% deoxycholic acid, 0.1% SDS, 2 mM MgCl_2_; Sigma-Aldrich) supplemented with proteases and phosphatases inhibitors and 50 mM 2- Chloroacetamide to inhibit the de-ubiquitinases. Lysates were centrifuged at 12000 rpm for 15 min at 4°C. Protein concentration was determined by pierce BCA protein assay kit and 500 µg of protein lysates were used as starting material for immunoprecipitation. Protein extracts were incubated overnight with 1 μg of anti-PDGFRβ (D-6) antibody and immunocomplexes recovered using 25 µL Protein G Sepharose beads (Sigma) for 1 h 30 min in the wheel at 4°C. The beads were washed with RIPA buffer and proteins were eluted in 2x Laemmli buffer supplemented with 1 mM DTT and boiled 5 min at 95°C. Western blots were performed as stated before and the membranes were incubated overnight at 4°C with the following primary antibodies and dilutions: anti-ubiquitin (1:1000) and anti-PDGFRβ (28E1). For analysis of whole cell lysates, 30 µg of protein lysates were used and membranes were incubated with anti - PDGFRβ (D-6) (1:500) and anti-GAPDH (1:1000) as loading control. The next day, blots were incubated 1 h at room temperature with goat anti-rabbit (1:10000), anti-mouse (1:5000) or anti-mouse light chain (1:1000) HRP-conjugated secondary antibodies and signal was detected as described above (see Western Blot analysis). Densitometric analysis was done with Image-J (NIH) software and ubiquitination levels were normalized to the corresponding levels of immunoprecipitated PDGFRβ.

### Confocal microscopy

Cells were cultured on collagen I pre-coated coverslips disposed on 24-well plates. 15000 fibroblasts per condition were cultured and they were serum-starved overnight prior to stimulation with 15 ng/mL PDGF-BB for 0, 5 and 10 min. After treatment, cells were fixed with parafolmaldehyde (PFA) 4% for 15 min and washed with PBS. Blocking and permeabilization was performed with 10% goat serum prepared in PBS with 0.25% Triton X-100 (Sigma-Aldrich). Cells were incubated with anti-PDGFRβ (28E1) (1:100) and anti-SNX1 (1:100) overnight at 4°C in a humid chamber. Coverslip were washed with PBS with 0.25% triton X-100 and followed by incubation with secondary antibodies Alexa Fluor 488 anti-rabbit and Alexa Fluor 594 anti-mouse (1:1000) together with Hoechst 33342 1 µM (14533; Sigma-Aldrich) for nuclei staining during 1h at room temperature. Coverslips were mounted in glass slides with ProLong Gold antifade reagent (P36930; ThermoFisher Scientific) and the edges were sealed with nail polish. Cells were visualized with inverted microscopy. Images of the specimens were collected with a TCS SPE-II confocal microscope (Leica Microsystems, Wetzlar, Germany), equipped with 63×/1.3 oil-immersion objective. Z-series images were obtained through the collection of serial, confocal sections at 0.4 μm intervals. To compare the confocal data, identical confocal settings were used for image acquisition of different experiments Quantitative co-localization analysis was performed with Imaris x64 v7.2.1 software (BITPLANE, Oxford Instruments), using object-based co-localization. For generation of SNX1 and PDGFRβ spots, particle size was measured and determined to 0.4 µm. The minimum distance between spots was set to 0.4 µm (distance from the center of spots), meaning that the spots has to be in touch for considering them as co-localizing particles. Finally, the number of co-localized spots were counted per cell.

### BrdU-based cell viability assay

Cells were cultures in 96-well plates and they were serum deprived overnight. Cells were treated with 25 ng/mL of PDGF-BB together with BrdU in low-serum DMEM. After 24 h incubation, the reaction was stopped and the incorporation of BrdU was detected with an HRP-secondary antibody, according to the manufacturer’s recommendations (BrdU Cell Proliferation ELISA Kit, Abcam).

### Cell migration assay

Normal and RDEB-derived fibroblasts were cultured in 96-well plates and they were grown until they reached full confluence. Cells were serum-starved overnight and the following day they were pre-incubated with 1 µg/mL mitomycin C (CAS 50-07-7, sc-3514A; Santa Cruz Biotechnologies) 1 h prior to the scratch generation to block proliferation. The scratch was performed with the the IncuCyte® 96 –well WoundMaker Tool (Essen Biosciences; Newark, UK). Cells were washed twice with low-serum DMEM and finally they were stimulated with vehicle or 5 ng/mL PDGF-BB in presence of 1µg/mL mitomycin C. The motility of cells was monitored during 30 h by time-lapse video microscopy using the Incucyte® Live Cell analysis system (Essen Biosciences). Quantification of wound closure was performed using the plugin “MRI-wound healing” developed for Image-J (NIH) software. In each experiment, wound closure data was normalized to the respective control, corresponding to the width at 0 h.

### Statistical analysis

Graph-Pad Prism 6 software (GraphPad, La Jolla, CA) was used for graphs generation and statistical analysis. Data are presented as mean ± s.e. (standard error) of the number of indicated experiments, unless otherwise mentioned. Appropriate statistical tests were performed based on the compared experimental conditions, including Student *t* test (computed with FDR ≤ 0.05 for correcting for multiple testing when necessary), one-way or two-way analysis of variance (ANOVA). Throughout the paper, statistically significant differences are indicated by : * *p* ≤ 0.05, ** *p* ≤ 0.01 and *** *p* ≤ 0.001.

## Data availability statement

The mass spectrometry proteomics data are publicly available and have been deposited to the ProteomeXchange Consortium via the PRIDE partner repository [48] with the dataset identifier PXD022985; reviewer account details: Username, Password: Ifuy3P8O

## Funding

This work was supported by the German Research Foundation (DFG) [grant number DE 1757/3-2] (JD), Swiss National Science Foundation [grant number 177088] (JD), the University and the Canton of Fribourg and was part of the SKINTEGRITY.CH collaborative research project.

### Abbreviations

NHF: normal human fibroblast
RDEB: recessive dystrophic epidermolysis bullosa
MS: mass spectrometry
LC: liquid chromatography
IP: immunoprecipitation
PDGFR: platelet-derived growth factor receptor
BrdU: bromodeoxyuridine
TGFβ: transforming growth factor β

